# A curated collection of transcriptome datasets to study the transcriptional response in blood and nasal samples following viral respiratory inoculation and vaccination

**DOI:** 10.1101/2024.09.23.614262

**Authors:** Zheng Lan, Tongli Zhang, Yu Zhang, Xuejun Sun, Chuwen Liu, Pei Liu, Muyao Tang, Meng Fu, James S. Hagood, Raymond J Pickles, Fei Zou, Xiaojing Zheng

**Affiliations:** Department of Biostatistics, University of North Carolina at Chapel Hill, Chapel Hill, North Carolina, USA; Industrial Engineering & Operations Research Department, College of Engineering, UC Berkeley, Berkeley, California, USA; Department of Probability and Statistics, School of Mathematical Sciences, University of Science and Technology of China, Hefei, Anhui, China; Department of Pediatrics, Division of Pulmonology, UNC School of Medicine, University of North Carolina at Chapel Hill, Chapel Hill, North Carolina, USA; Marsico Lung Institute, UNC School of Medicine, University of North Carolina at Chapel Hill, Chapel Hill, North Carolina, USA; Department of Microbiology & Immunology, School of Medicine, University of North Carolina at Chapel Hill, Chapel Hill, North Carolina, USA; Department of Pediatrics, Division of Infectious Diseases, UNC School of Medicine, University of North Carolina at Chapel Hill, Chapel Hill, North Carolina, USA

**Keywords:** Transcriptomics, Bioinformatics, Vaccination, Inoculation, Respiratory viral infection, Influenza viruses, COVID, Respiratory syncytial viruses (RSV), Human rhinoviruses (HRV) Whole Blood, PBMC

## Abstract

Our understanding of the human immune system’s response to viral respiratory tract infections (VRTIs) and vaccines, including the molecular mechanisms and correlates of protection, remains incomplete. Extensive transcriptomic data from inoculation and vaccination studies have been deposited in publicly available databases. However, these studies are often separate and difficult to locate. We provide a curated compendium of public gene expression data repositories for researchers to reanalyze transcriptomes from human whole blood, peripheral blood mononuclear cells (PBMCs), and nasal swab samples. This enables the study of transcriptional responses to viral inoculation or vaccination. This collection includes 18 datasets from inoculation studies and 37 datasets from vaccination studies, sourced from the NCBI Gene Expression Omnibus (GEO), ImmPort Shared Data, and ArrayExpress.

## Introduction

Viral respiratory tract infections (VRTIs) are widespread globally, causing significant illnesses and hospitalizations each year.^1^ The importance of studying VRTIs has grown in light of the global COVID-19 pandemic.^2^ VRTIs involve a range of viruses, including respiratory syncytial viruses (RSV), human rhinoviruses (HRV), and influenza and coronaviruses.^3^ Most VRTIs lack both effective antiviral therapies and approved vaccines.^4^

In-depth investigation of human immune responses to inoculation and vaccination is critically needed. To facilitate further research, we have compiled a curated dataset collection, which includes transcriptomic data from the blood and nasal swabs of volunteers who underwent inoculation and vaccination, specifically for influenza, COVID-19, RSV, and HRV.

Our curated dataset collection is divided into two categories: inoculation studies and vaccination studies. The inoculation collection includes 18 datasets with 429 participants, while the vaccination collection comprises 37 datasets with 2,084 participants. All datasets were sourced from the NCBI Gene Expression Omnibus (GEO), ImmPort Shared Data, and the ArrayExpress Collection at EMBL-EBI.^5–7^ These datasets have been organized into a publicly accessible format for download and further analysis.

## Methods

The datasets were gathered using specific search queries across GEO, ArrayExpress, ImmPort, and Google search. The queries for inoculation studies included terms related to humans, blood, PBMCs, respiratory challenges, and viral inoculations. The vaccination study queries followed similar criteria, focusing on humans, blood, PBMCs, transcriptomes, and vaccines or vaccinations. The search queries used for GEO, ArrayExpress, and Google search to find inoculation studies are as follows:

### GEO and ArrayExpress Inoculation Search Query

(“humans”[MeSH Terms] OR “Homo sapiens”[Organism] OR Homo sapiens[All Fields]) AND ((“blood”[Subheading] OR “blood”[MeSH Terms] OR blood[All Fields]) OR PBMC[All Fields] OR PBMCs[All Fields]) AND respiratory[All Fields] AND (challenge[All Fields] OR “experimentally infected”[All Fields] OR inoculated[All Fields])

### Google Inoculation Search Query

Homo sapiens AND (blood OR PBMC OR PBMCs) AND transcriptome AND respiratory AND (inoculation OR challenge OR “experimentally infected” OR inoculated)

The search queries used for GEO, ArrayExpress, and Google to find vaccination studies are as follows:

### GEO and ArrayExpress Vaccine Search Query

(“humans”[MeSH Terms] OR “Homo sapiens”[Organism] OR Homo sapiens[All Fields]) AND ((“blood”[Subheading] OR “blood”[MeSH Terms] OR blood[All Fields]) OR PBMC[All Fields] OR PBMCs[All Fields]) AND respiratory[All Fields] AND ((“vaccination”[MeSH Terms] OR inoculation[All Fields]) OR (“vaccines”[MeSH Terms] OR vaccine[All Fields]) OR (“vaccines”[MeSH Terms] OR vaccines[All Fields]) OR (“vaccination”[MeSH Terms] OR vaccination[All Fields]))

### Google Vaccine Search Query

Homo sapiens AND (blood OR PBMC OR PBMCs) AND transcriptome AND respiratory AND (vaccine OR vaccines OR vaccination)

For **ImmPort**, vaccine studies were located using the following filters:

- Species: Homo sapiens
- Research Focus: Vaccine Response
- Condition or Disease: COVID-19 AND Influenza

## Results

The majority of datasets obtained were derived from human blood, PBMCs, and nasal swabs, generated using Illumina or Affymetrix platforms or through RNA sequencing. Every dataset identified through this search was carefully curated manually. This involved thoroughly reviewing the dataset descriptions, examining the study designs, and reading through the related original articles on PubMed. Ultimately, we only included studies that involved human whole blood, PBMCs, and nasal swabs linked to VRTI inoculation or vaccination for our dataset collection. According to these criteria, we retained 18 inoculation datasets and 37 vaccine-related datasets.

The data selection process for inoculation is summarized in **Figure 1a**. Using GEO and ArrayExpress inoculation query, 108 studies were found with Homo sapiens as the primary organism. Google search returned 92 inoculation studies. The data selection process for vaccination is summarized in **Figure 1b**. GEO and ArrayExpress Vaccine query identified 82 studies where Homo sapiens is the top organism. Google search produced 89 vaccination studies. ImmPort search yielded 99 vaccination studies.

**Figure 1.**
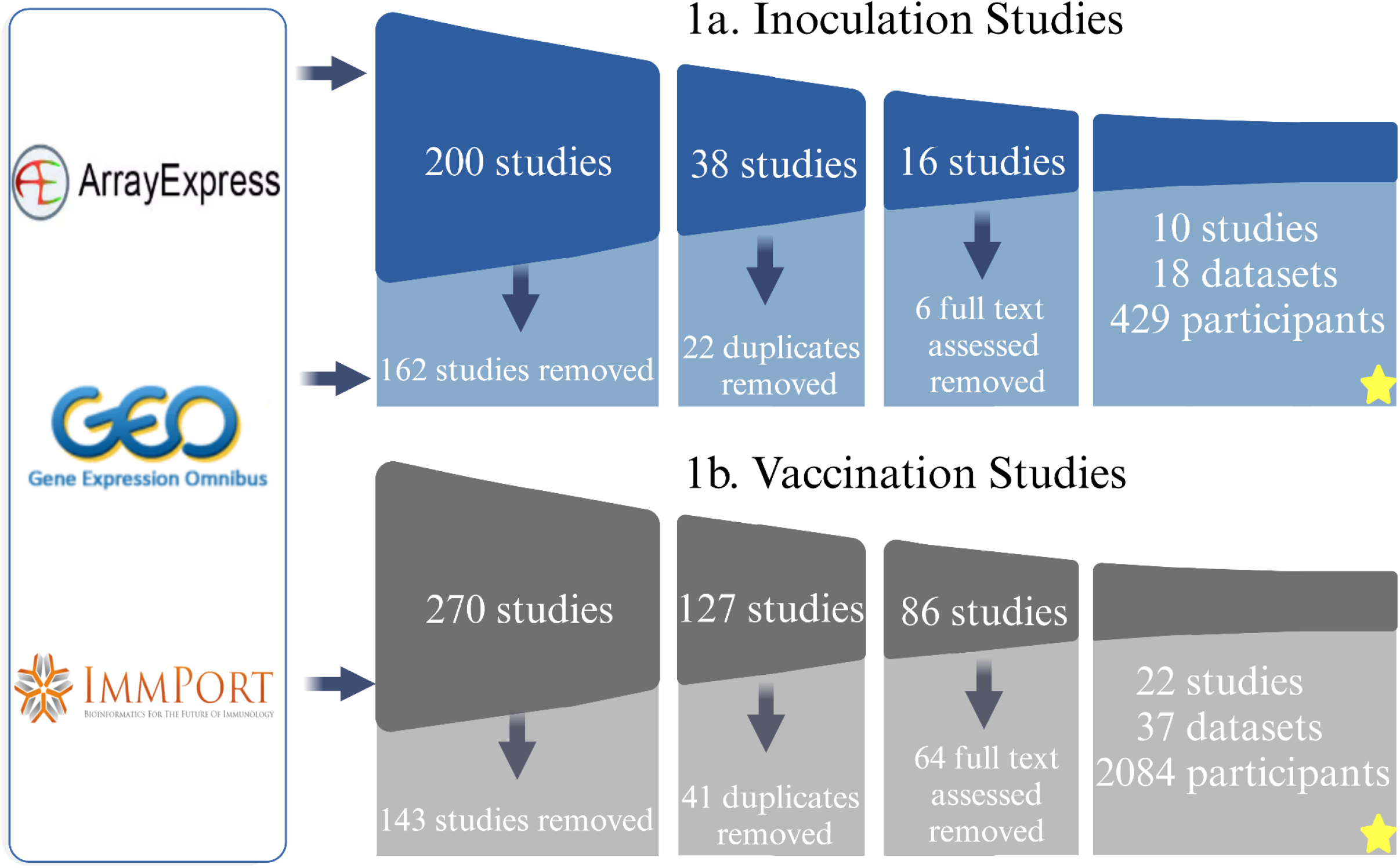
Flowchart depicting the collection, filtering, and curation steps leading to the construction of the inoculation (Fig. 1a) and vaccination (Fig. 1b) datasets compendium, respectively.

Figure 2. illustrates the distribution of participants over time in inoculation studies across different viruses, with a notable peak of over 400 participants on day 0. Notably, 56 participants had both blood and nasal samples and they were counted separately. Subsequent time points show smaller but consistent sample collections, with varying contributions from different pathogens. H1N1, H3N2, and HRV are the most frequently sampled pathogens, while SARS-CoV-2 and RSV contribute fewer samples to the dataset.

**Figure 2.**
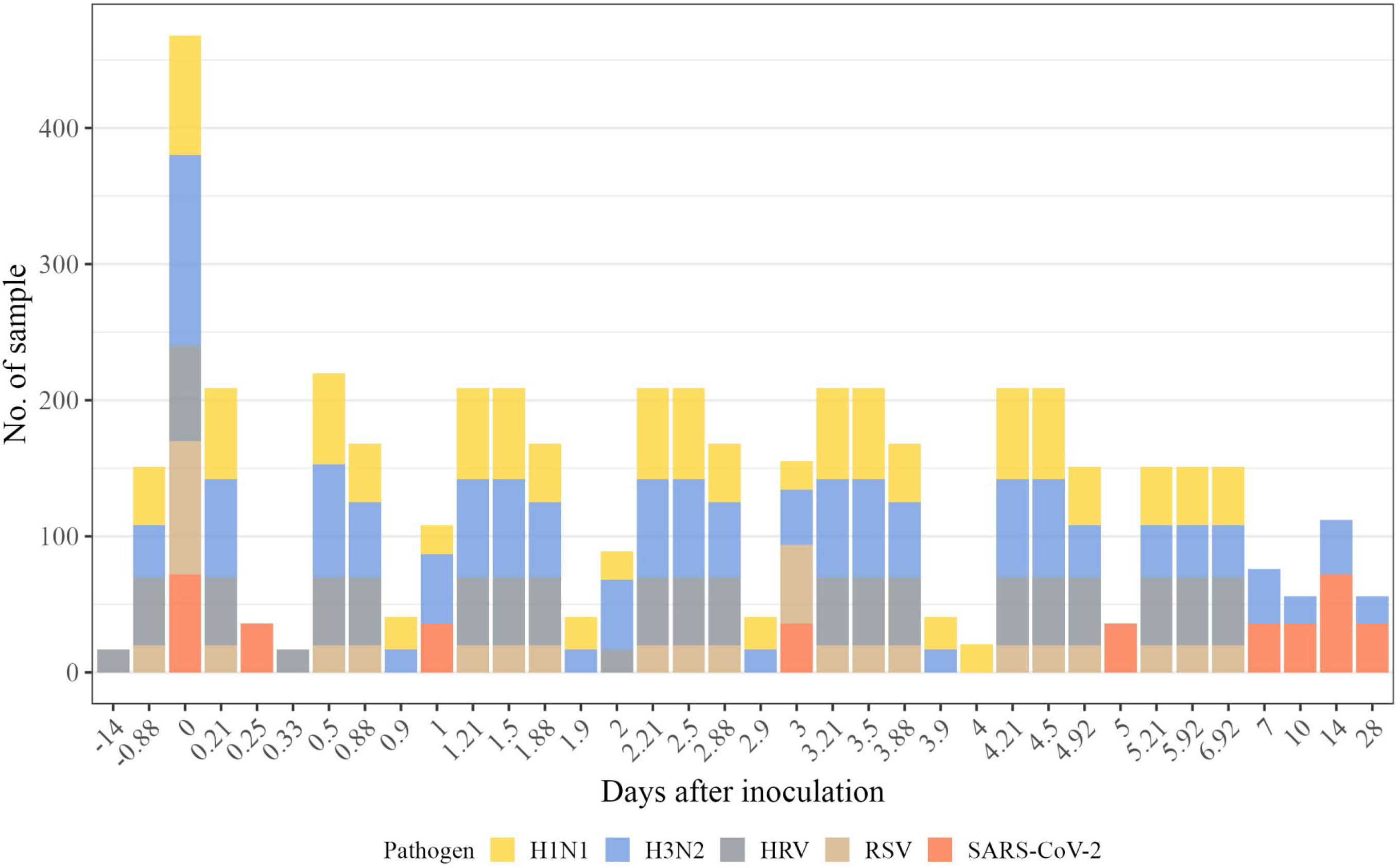
A collection of transcriptome datasets of transcriptional responses to inoculation over time across varying viruses.

Figure 3. summarizes the distribution of participants over time in vaccination studies, covering COVID-19 and influenza vaccines and their types. The peak occurred on day 0, with over 2,000 participants. Participant recruitment was enriched during the first week.

**Figure 3.**
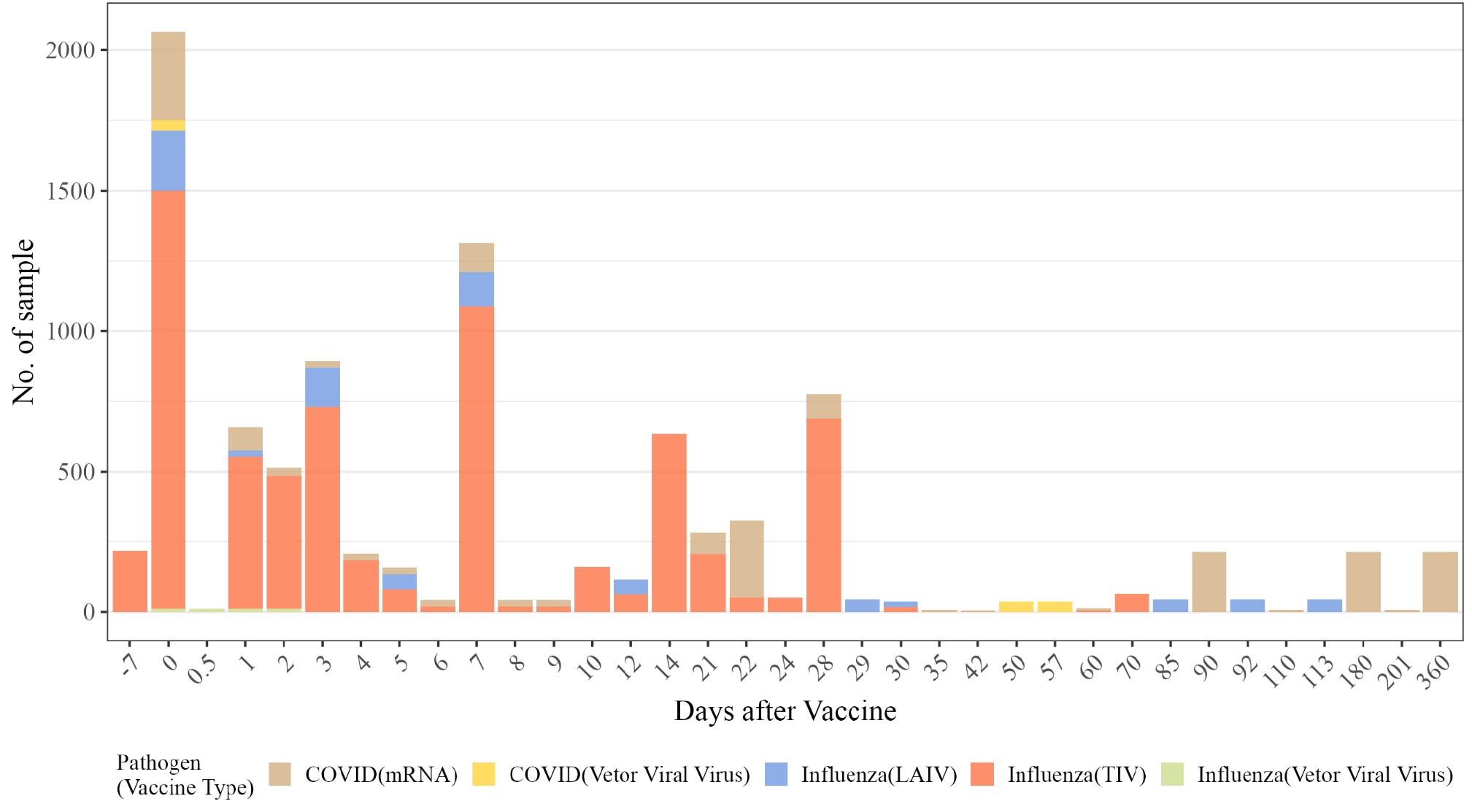
A collection of transcriptome datasets of transcriptional responses to vaccination over time covering COVID-19 and influenza vaccines and their types.

The complete list of inoculation datasets included in our collection can be found in **Table 1**.

**Table 1.** Complete list of inoculation datasets.

| Pathogen | Sample Type | Sample Size | Profiling Platform | Time Point | Accession Number | Citation |
| --- | --- | --- | --- | --- | --- | --- |
| HRV | Peripheral blood | 50 | Microarray(Affymetrix ) | -0.88, 0, 0.21, 0.5, 0.88, 1.21, 1.5, 1.88, 2.21, 2.5, 2.88, 3.21, 3.5, 3.88, 4.21, 4.5, 4.92, 5.21, 5.92, 6.92 | GSE73072 | 8 |
| RSV |  | 20 |  |  |  |  |
| H3N2 |  | 38 |  |  |  |  |
| H1N1 |  | 43 |  |  |  |  |
| HRV | Peripheral blood | 20 | Microarray(Affymetrix ) | 0 | GSE17156 | 9 |
| RSV |  | 20 |  |  |  |  |
| H3N2 |  | 17 |  |  |  |  |
| H3N2 | Peripheral blood | 17 | Microarray(Affymetrix ) | 0, 0.21, 0.50, 0.88, 1.21, 1.50, 1.88, 2.21, 2.50, 2.88, 3.21, 3.50, 3.88, 4.21, 4.50 | GSE30550 | 10 |
| H3N2 | Whole blood | 11 | Microarray(Illumina) | 0,0.50,1,2 | GSE61754 | 11 |
| SARS-CoV-2 | Blood and nose swabs | 36 | RNA-seq | 0, 0.25, 14, 28(blood)<br>0,1,3,5,7,10,14(nose swabs) | E-MTAB-12993 | 12 |
| H3N2 | Blood and nose swabs | 20 | RNA-seq | 0, 1, 2, 3, 7, 10, 14, and 28(blood)<br>0,1, 2,3,7,14(nose swabs) | E-MTAB-13038<br>E-MTAB-13041 |  |
| HRV | nose swabs | 17 | Microarray(Affymetrix) | -14,0.33, 2.00 | GSE11348 | 13 |
| H1N1 | Whole blood | 21 | Microarray(Illumina) | 0, 1, 2, 3, 4 | GSE90732 | 14 |
| RSV | nose swabs | 58 | RNA-seq | 0,3 | GSE155237 | 15 |
| H1N1 | PBMC | 24 | Microarray(Affymetrix ) | 0, 0.21, 0.50, 0.90, 1.21, 1.50, 1.90, 2.21, 2.50, 2.90, 3.21, 3.50, 3.90, 4.21, 4.50 | GSE52428 | 16 |
| H3N2 |  | 17 |  |  |  |  |

PBMC: peripheral blood mononuclear cells
Time point: Unit is day. 0 is the baseline, negative number means days before inoculation, and positive means days after inoculation.
Accession number: datasets from GEO start with "GSE", datasets from ArrayExpress start with "E-MTAB"

We have collected transcriptomes from 18 cohorts across 10 inoculation studies, totaling 429 participants. This collection covers inoculations of five respiratory viruses: H1N1, H3N2, RSV, HRV, and SARS-CoV-2. The time span ranges from -14 to 28 days before and post for inoculation. The sample source includes blood (n=14) and nasal swabs (n=4). The transcriptomic data types include microarrays from Illumina (n=2) and Affy (n=11); bulk RNA-seq (n=5) **(Figure 4)**.

**Figure 4.**
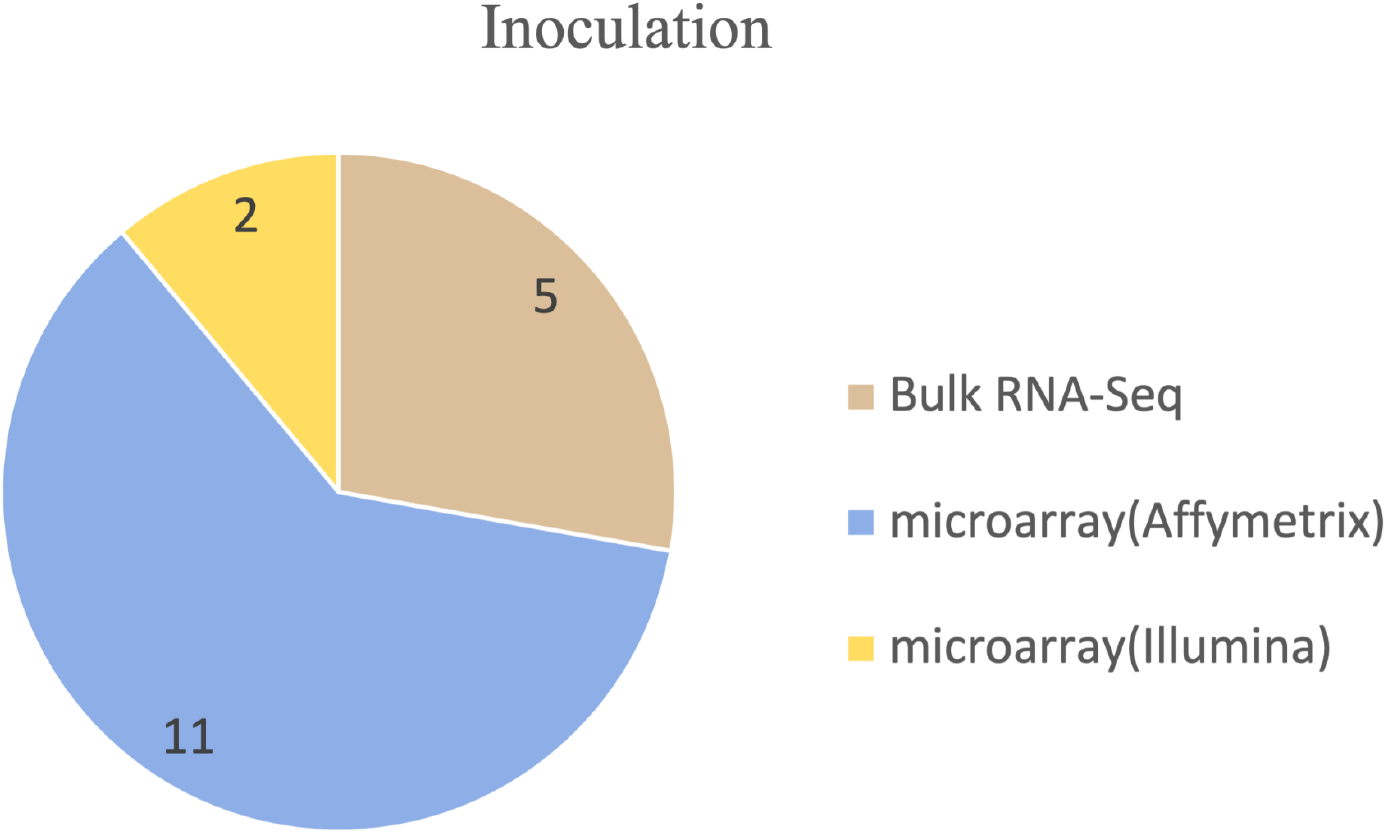
Distribution of Inoculation Datasets Across Different Profiling Platforms.

The complete list of vaccination datasets included in our collection can be found in **Table 2**.

**Table 2.**
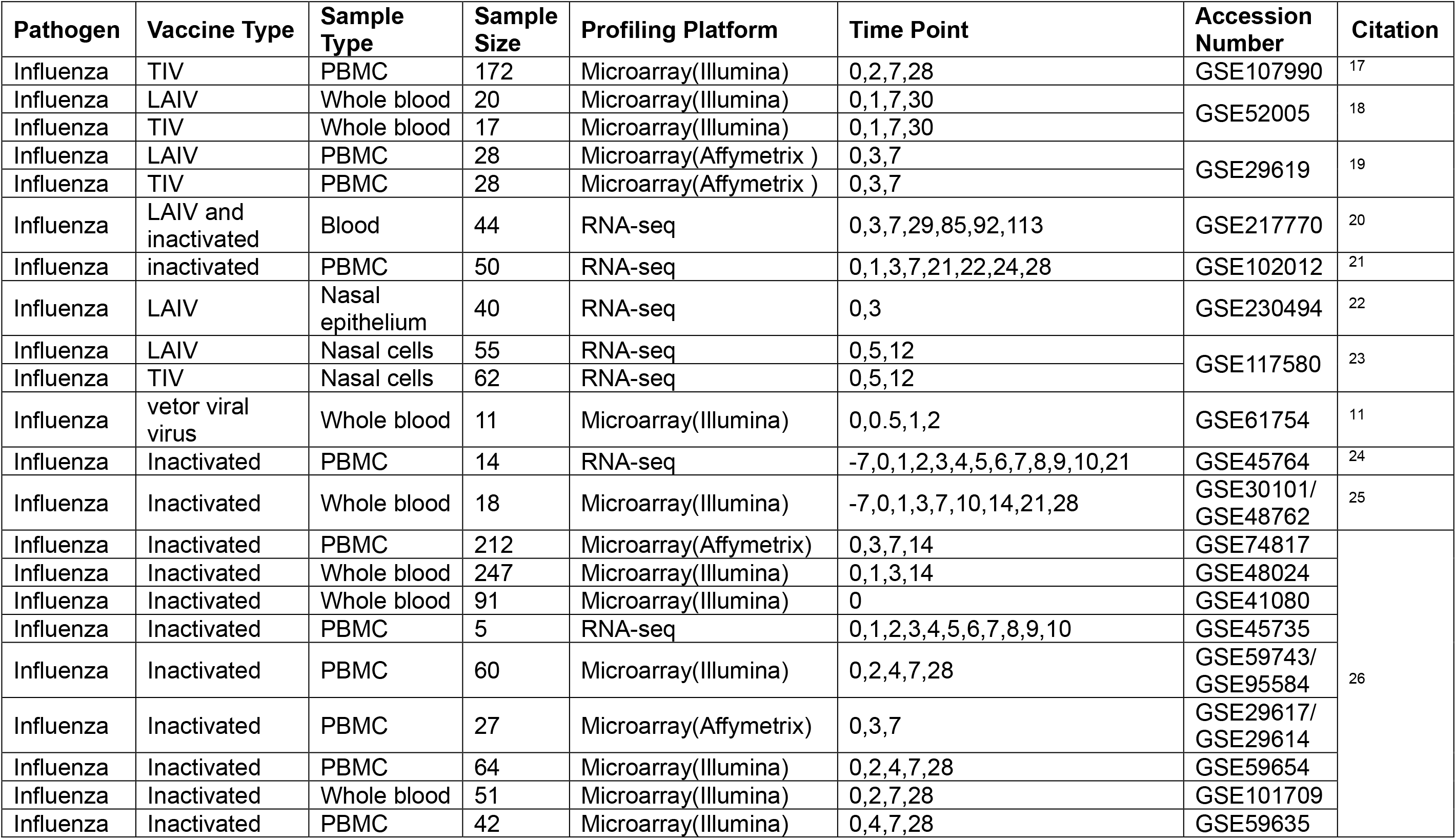
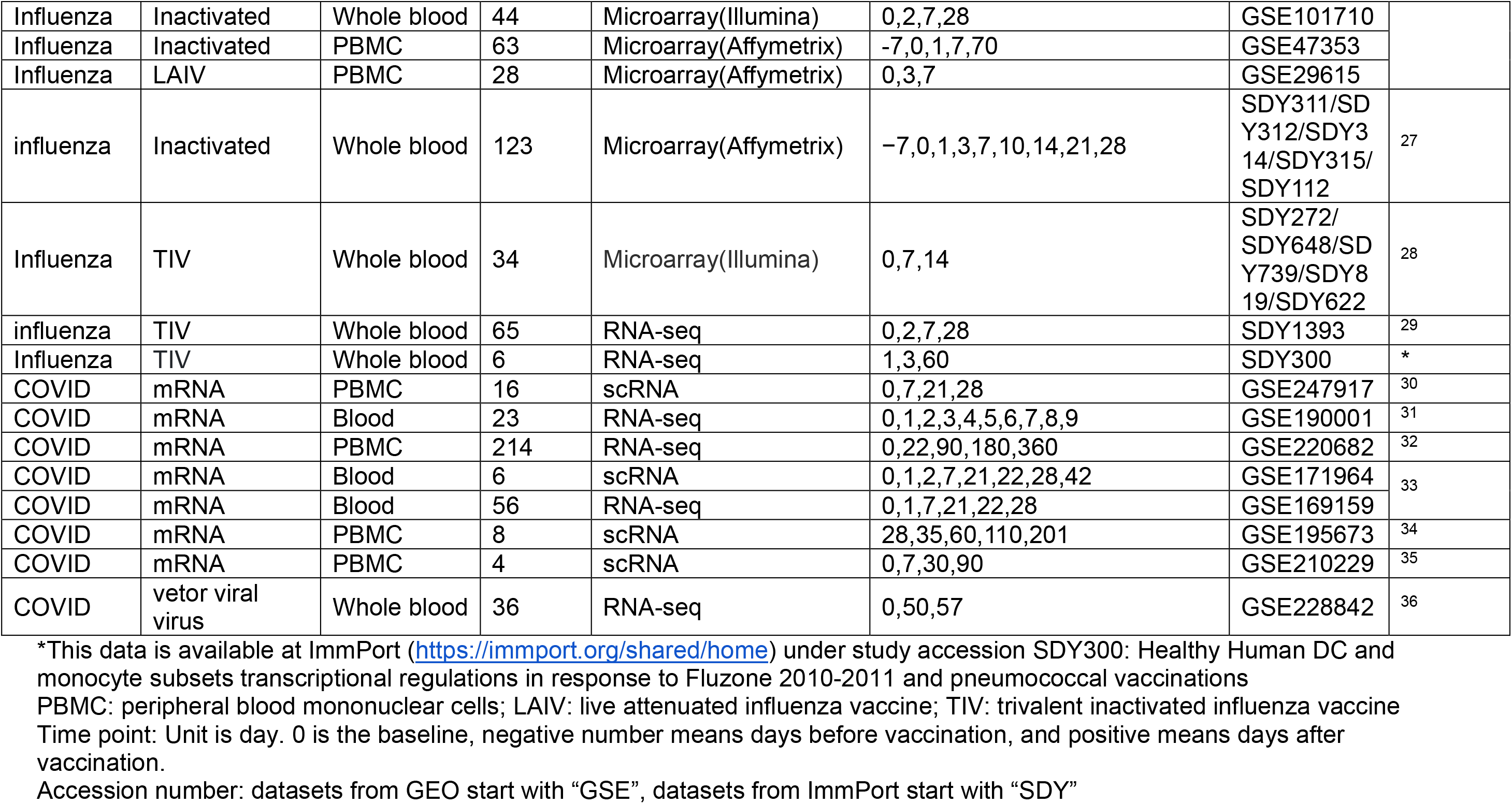
Complete list of vaccine datasets.

We have gathered transcriptomes from 37 cohorts across 22 vaccine studies, totaling 2084 participants. This collection focuses on studying Influenza and COVID-19 vaccines. The time span ranges from -28 to 360 days. The data source includes blood (n=34) and nasal swabs (n=3). The transcriptomic data types include microarrays from Illumina (n=13) and Affy (n=7); bulk RNA-seq (n=13) and scRNA-seq (n=4) **(Figure 5)**. Vaccine types: TIV (n=22) and LAIV (n=6) were used for influenza vaccination, mRNA vaccine (n=7) was used for COVID-19 vaccination, and vector adenovirus (n=2) was used for both influenza and COVID-19, one of each.

**Figure 5.**
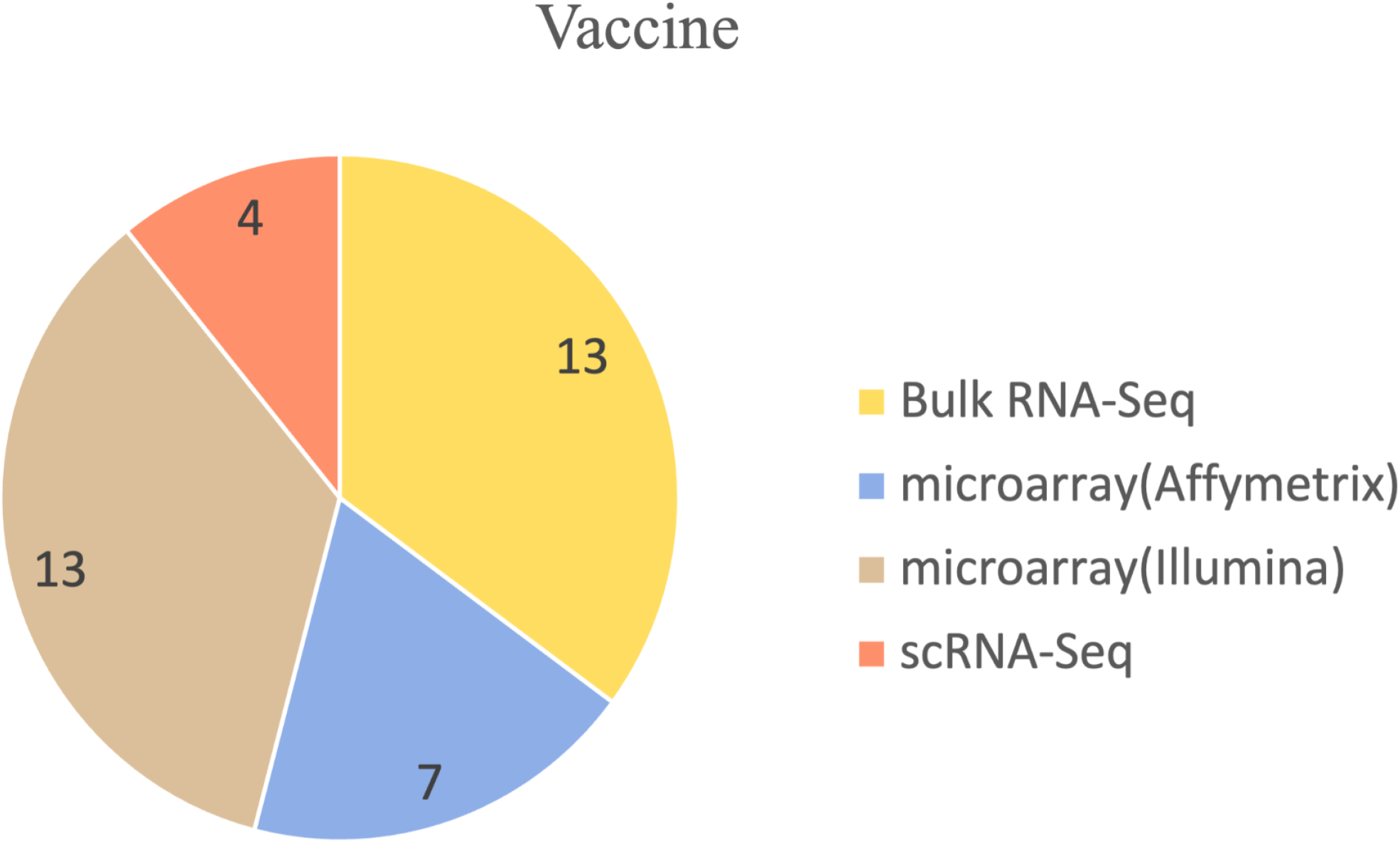
Distribution of Vaccine Datasets Across Different Profiling Platforms.

Histogram of the time points pre- and post-inoculation available in our compendium. Each virus is indicated by a different color. The height of the bars represents the number of participants with available gene expression data.

Histogram of the time points pre- and post-vaccination is available in our compendium. Each vaccine and type is indicated by a different color. The height of the bars represents the number of participants with available gene expression data.

## Data availability

All datasets in our curated collection are also accessible to the public on the NCBI GEO website at https://www.ncbi.nlm.nih.gov/gds/ ArrayExpress at https://www.ebi.ac.uk/biostudies/arrayexpress/studies and ImmPort at https://immport.org/shared/home. Each dataset is cited in the manuscript using its GEO number, ArrayExpress number, or ImmPort study number.

## Acknowledgment

We extend our gratitude to all the researchers who chose to share their datasets publicly by depositing them in GEO, ArrayExpress, and ImmPort.

